# If Metformin Inhibited the Mitochondrial Glycerol Phosphate Dehydrogenase It Might Not Benefit Diabetes

**DOI:** 10.1101/2020.03.28.013334

**Authors:** Michael J. MacDonald, Israr-ul H. Ansari, Melissa J. Longacre, Scott W. Stoker

## Abstract

The mitochondrial glycerol phosphate dehydrogenase is the rate-limiting enzyme of the glycerol phosphate shuttle. It was recently claimed that metformin, a first line drug used worldwide for the treatment of type 2 diabetes, works by inhibiting the mitochondrial glycerol phosphate dehydrogenase 30-50% thus suppressing hepatic gluconeogenesis. This enzyme is 30-60 fold higher in the pancreatic islet than in liver. If metformin actually inhibited the enzyme, why would it not inhibit insulin secretion and exacerbate diabetes? Total body knockout of the mitochondrial glycerol phosphate dehydrogenase does not inhibit insulin secretion because insulin cells and liver cells possess the malate aspartate shuttle that is redundant to the action of the glycerol phosphate shuttle. In view of these and other apparent inconsistencies we reassessed the idea that metformin inhibited the mitochondrial glycerol phosphate dehydrogenase. We measured the enzyme’s activity in whole cell homogenates and mitochondria of insulin cells and liver cells using four different enzyme assays and were unable to show that metformin directly inhibits the enzyme. We conclude that metformin does not actually inhibit the enzyme. If it did, it might not be efficacious as a diabetes medicine.

---

The mitochondrial glycerol phosphate dehydrogenase is the rate-limiting enzyme of the glycerol phosphate shuttle. It was recently claimed that metformin, a first line drug used worldwide for the treatment of type 2 diabetes, works by inhibiting the mitochondrial glycerol phosphate dehydrogenase 30-50% thus suppressing hepatic gluconeogenesis^1^. This enzyme is 30-60 fold higher in the pancreatic islet than in liver^2^. If metformin actually inhibited the enzyme, why would it not inhibit insulin secretion and exacerbate diabetes? Total body knockout of the mitochondrial glycerol phosphate dehydrogenase does not inhibit insulin secretion^3,4^ because insulin cells and liver cells possess the malate aspartate shuttle that is redundant to the action of the glycerol phosphate shuttle^5,6,7^. In view of these and other apparent inconsistencies we reassessed the idea that metformin inhibited the mitochondrial glycerol phosphate dehydrogenase. We measured the enzyme’s activity in whole cell homogenates and mitochondria of insulin cells and liver cells using four different enzyme assays and were unable to show that metformin directly inhibits the enzyme. We conclude that metformin does not actually inhibit the enzyme. If it did, it might not be efficacious as a diabetes medicine.

Metformin is a first line drug for the treatment of type 2 diabetes. After decades of use in millions of patients and many biomolecular studies of metformin, the exact mechanism of metformin’s antihyperglycemic action is still not completely understood. Metformin’s blood glucose lowering effect is through its decreasing hepatic glucose production. In the year 2000 metformin, at higher than clinically achievable concentrations, was shown to decrease cellular respiration through specific inhibition of complex I of the mitochondrial electron transport chain^8^. Shortly thereafter metformin was shown to activate AMP-activated protein kinase (AMPK) which suppresses gluconeogenesis in the liver by increasing the phosphorylation of the AMPK catalytic α subunit^9,10^. Liver Kinase B1 (LKB1) is the upstream kinase that catalyzes the phosphorylation of threonine 172 in the α subunit of AMPK. In support of this idea, metformin fails to improve hyperglycemia in mice with the liver-specific knockout of the kinase LKB1^11^. Although both the inhibition of mitochondrial complex I and activation of AMPK are still believed to be possible explanations for metformin’s blood glucose-lowering effect their actions are each believed to be indirect. Therefore, the search for metformin’s direct target is still active.

Madiraju et al^1^ recently claimed that metformin’s blood glucose-lowering mechanism is through a 30-50% direct inhibition of the mitochondrial glycerol phosphate dehydrogenase (mGPD) suppressing gluconeogenesis in liver^1^. mGPD is imbedded in the outer surface of the inner mitochondrial membrane freely accessible to cytosolic metabolites. mGPD and the cytosolic glycerol phosphate dehydrogenase (cGPD) catalyze the two enzyme reactions of the glycerol phosphate shuttle, a redox shuttle that via the mGPD reaction utilizes the mitochondrial electron transport chain to oxidize glycerol phosphate to dihydroxyacetone phosphate. The cGPD then uses dihydroxyacetone phosphate to reoxidize cytosolic NADH to NAD^+^ to allow cytosolic pathways that require NAD^+^ to function normally. Unlike the cGPD reaction that is reversible, the mGPD reaction is unidirectional and keeps the glycerol phosphate shuttle operating only in the direction of converting NADH to NAD^+^ in the cytosol of the cell (See Fig. 2 in Ref. 7). At first glance inhibition of mGPD might be expected to increase the liver cytosolic NADH/NAD^+^ ratio which would appear to be consistent with metformin’s proposed mechanism of action. Prior to the introduction of metformin to the market as an antidiabetic medication, its precursor phenformin which is a similar but a stronger biguanide medication, was withdrawn from the market due to its causing numerous severe cases of lactic acidosis in diabetes patients. An increased blood lactate/pyruvate ratio is consistent with the idea that the biguanide family of medications might inhibit reoxidation of NADH. Although metformin was also feared to be a primary risk factor for lactic acidosis, a recent study of a large number of patients showed that there is little or no risk of lactic acidosis associated with proper dosing of metformin in the absence of complicating factors such as renal or heart disease^12^.

Somewhat consistent with the idea that metformin might increase the blood’s NADH/NAD^+^ ratio Madiraju et al^1^ showed that metformin slightly increased the plasma and liver lactate/pyruvate ratio in rats. They found that metformin had no effect on plasma glucose in the mGPD knockout mouse^1^ that was a gift from our laboratory^3^. Although they attributed this absence of an effect as more evidence that metformin acted through mGPD, this result is subject to other interpretations. Also consistent with a mechanism of metformin increasing the NADH/NAD^+^ ratio, Madiraju et al^13^ showed in a later study of rats that metformin decreased gluconeogenesis from the reduced substrates lactate and glycerol, but not from the oxidized substrates pyruvate and alanine. Many of these data are puzzling. The mGPD knockout mouse possesses absolutely no mGPD in any tissue^3^. In contradiction to the idea of metformin significantly influencing the NADH/NAD^+^ ratio via inhibition of mGPD we previously showed that in this mGPD knockout mouse the level of metabolites in liver including the cytosolic lactate/pyruvate and glycerol phosphate/dihydroxyacetone phosphate ratios that each reflect the cytosolic NADH/NAD^+^ ratio are normal^3^. The reason these ratios are normal in the mGPD knockout mouse is because liver cells possess another mitochondrial NADH/NAD^+^ shuttle called the malate aspartate shuttle^5,6,7^. The action of this shuttle is redundant to the action of the glycerol phosphate shuttle.

The idea that metformin’s antihyperglycemic action involves inhibition of hepatic mGPD is also puzzling because the enzyme activity of mGPD in pancreatic islets and pancreatic beta cells is known to be 30-60 fold higher than in liver^2^. If it were not for the redundancy of the malate aspartate NAD^+^/NADH shuttle in the insulin cell, with the glycerol phosphate shuttle in beta cells^6,7^ (See Fig. 4 in Ref. 7), inhibition of mGPD by metformin might be expected to have a diabetogenic effect by inhibiting insulin secretion rather than an antidiabetogenic effect via the liver. In support of the idea of the known redundancy of the two shuttles, insulin secretion from pancreatic islets isolated from the mGPD knockout mouse is not inhibited unless the malate aspartate shuttle is also inhibited^4^. The malate aspartate shuttle is very active in the liver relative to the glycerol phosphate shuttle, so why would inhibition of the glycerol phosphate shuttle alone in the liver influence glucose production? If there was not a shuttle redundant to the glycerol phosphate shuttle and if metformin did indeed inhibit mGPD, it might not be beneficial to diabetes. Because of these inconsistencies, we used four different mGPD enzyme assays to reassess the effect of metformin on mGPD activity. We used two mGPD assays^2,14,15^ that we have used numerous times previously including when we found that mGPD activity is much higher in pancreatic islets than in liver^2^. We also used the mGPD assay used by Madiraju et al^1^, as well as one other mGPD assay^16^.

First, we studied the effect of metformin and phenformin on mGPD using iodonitrotetrazolium blue (INT) as an electron acceptor^2,14^ in a timed-and-stopped mGPD enzyme assay that we have previously used for numerous publications (> 10). We measured the effect of concentrations of metformin between 10 μM and 1 mM using whole cell homogenates and isolated mitochondria from both liver and a much richer source of mGPD (the INS-1 832/13 insulinoma cell line (pure beta cells)). We also tested the effect of phenformin on mGPD enzyme activity using concentrations of phenformin between 250 μM and 10 mM. Neither of these drugs inhibited mGPD whatsoever (Table 1).

**Table 1.** No inhibition of the mitochondrial glycerol phosphate dehydrogenase (mGPD) by metformin or phenformin. Electron acceptor: iodonitrotetrazolium blue.

| **Metformin Experiments** |  |  |  |  |
| --- | --- | --- | --- | --- |
| **Enzyme Source** | INS-1 832/13 mitochondria | INS-1 832/13 homogenate | Liver mitochondria | Liver homogenate |
| **Control Specific mGPD Activity**<br>(nmol dye reduced/min/mg protein (N)) | 164 ± 4.6 (8) | 39 ± 0.71 (6) | 8.3 ± 0.6 (16) | 1.2 ± 0.07 (8) |
| **Concentration of Metformin** | **mGPD Activity (% of Control (mean ± SE (N)))** |  |  |  |
| **None** (control) | 100 ± 2.5 (8) | 100 ± 0.7 (6) | 100 ± 1.5 (16) | 100 ± 5.5 (8) |
| **10 µM** | 104 ± 0.9 (4) | 99 ± 2.5 (6) | 98 ± 4.4 (12) | 95 ± 6.3 (8) |
| **50 µM** | 102 ± 3.7 (4) | 102 ± 1.6 (6) | 99 ± 2.8 (8) | 106 ± 5.3 (8) |
| **100 µM** | 104 ± 3.5 (4) | 101 ± 1.8 (6) | 99 ± 2.4 (4) | 116 ± 4.6 (8) |
| **250 µM** | 104 ± 2.7 (4) | 107 ± 0.6 (4) | 98 ± 4.4 (4) | 110 ± 4.0 (4) |
| **0.5 mM** | 101 ± 2.5 (4) | 101 ± 1.0 (6) | 104 ± 1.6 (8) | 115 ± 6.0 (4) |
| **1 mM** | 97 ± 1.2 (4) | 102 ± 0.6 (6) | 109 ± 5.5 (4) | 121 ± 5.7 (4) |
| **Phenformin Experiments** |  |  |  |  |
| **Enzyme Source** | INS-1 832/13 mitochondria | INS-1 832/13 homogenate | Liver mitochondria | Liver homogenate |
| **Control Specific mGPD Activity**<br>(nmol dye reduced/min/mg protein (N)) | 194 ± 4.7 (4) | 34 ± 0.97 (4) | 6.6 ± 0.15 (4) | 1.6 ± 0.64 (10) |
| **Concentration of Phenformin** | **mGPD Activity (% of Control (mean ± SE (N)))** |  |  |  |
| **None** (control) | 100 ± 2.4 (4) | 100 ± 1.0 (8) | 100 ± 2.3 (4) | 100 ± 3.6 (10) |
| **250 µM** | 93 ± 2.3 (4) | 100 ± 1.5 (4) | 101 ± 0.5 (4) | 111 ± 6.8 (4) |
| **0.5 mM** | 94 ± 2.1 (4) | 98 ± 1.6 (4) | 100 ± 2.7 (4) | 110 ± 2.1 (5) |
| **1 mM** | 92 ± 2.4 (4) | 100 ± 2.4 (4) | 103 ± 1.8 (4) | 99 ± 4.1 (6) |
| **10 mM** | 89 ± 2.2 (4) | 95 ± 1.5 (6) | 109 ± 1.4 (3) | 105 ± 3.0 (4) |

We also tested the effect of metformin on mGPD enzyme activity with three other enzyme assays employing different electron acceptors^1,2,15,16^. We tested the effect of 250 μM metformin on the mitochondria isolated from the INS-1 832/13 cell line and the human liver cell line Huh 7.5. We measured mGPD enzyme activity in continuous spectrophotometric assays with the electron acceptor DCIP that we also used previously^2,15^, as well as with cytochrome c as an electron acceptor according to the method used by Madiraju et al^1^ (in which they observed inhibition of mGPD), and also with cytochrome c according to a method used by Rauchová et al^16^. With each of the three additional mGPD enzyme assays no inhibition of mGPD activity was observed (Table 2).

**Table 2.** No inhibition of mGPD enzyme activity by metformin in three different enzyme assays when using cytochrome c or DCIP as an electron acceptor.

| Enzyme Source | Concentration of Metformin |  |
| --- | --- | --- |
| | None<br>(Control) | 250 μM |
| | mGPD activity (% of Control (mean ± SE (N))) | |
| INS-1 832/13 mitochondria | 100 ± 5.9 (4)<sup>a</sup> | 102 ± 1.3 (4)<sup>a</sup> |
| INS-1 832/13 mitochondria | 100 ± 2.2 (4)<sup>b</sup> | 110 ± 3.3 (4)<sup>b</sup> |
| INS-1 832/13 mitochondria | 100 ± 0.6 (4)<sup>c</sup> | 112 ± 1.5 (4)<sup>c</sup> |
| Huh 7.5 Liver cell mitochondria | 100 ± 3.1 (4)<sup>a</sup> | 103 ± 2.2 (4)<sup>a</sup> |
| Huh 7.5 Liver cell mitochondria | 100 ± 6.4 (4)<sup>b</sup> | 106 ± 1.9 (4)<sup>b</sup> |
| Huh 7.5 Liver cell mitochondria | 100 ± 3.6 (8)<sup>c</sup> | 105 ± 3.6 (8)<sup>c</sup> |

<sup>a</sup>Electron receptor: cytochrome c<sup>1</sup>
<sup>b</sup>Electron receptor: cytochrome c<sup>16</sup>
<sup>c</sup>Electron receptor: DCIP<sup>2,15</sup>

We conclude from our data that metformin’s property of improving hyperglycemia does not involve direct inhibition of mGPD. Since the activity of mGPD is 30-60 fold higher in the insulin-producing pancreatic beta cell than in liver^2^ (Table 1). If metformin actually did inhibit mGPD and if it were not for the malate aspartate shuttle redundant to the action of the glycerol phosphate, metformin might not be efficacious as a diabetes medicine.

## METHODS

mGPD enzyme activity was measured with four different enzyme assays with different electron acceptors. When mGPD enzyme activity (Table 1) was measured according to the method of Gardner^2,14^, the enzyme reaction mixture contained 50 mM D-L-glycerol phosphate as the D-L mixture, 1 mM KCN, 50 mM Bicine buffer, pH 8.0, and 4 mM iodonitrotetrazolium violet and enzyme source incubated at 37° for 30 min. The enzyme reaction was stopped by adding ethyl acetate. The reaction mixture was centrifuged briefly and the ethyl acetate phase of the mixture containing the reduced iodonitrotetrazolium dye was removed and its absorbance measured at 490 nm. Absorbance values from reaction mixtures containing all ingredients except glycerol phosphate were subtracted from reaction mixtures containing all ingredients to give the enzyme rate attributable to mGPD activity.

mGPD enzyme activity was also measured in a continuous spectrophotometric assay at 600 nm at 38° in a reaction mixture containing 50 mM potassium phosphate buffer, pH 7.6, 1 mM KCN and 47.5 μM 2,6-dichlorophenolindophenol (DCIP) with or without 25 mM L-glycerol phosphate as the DL mixture as previously described^2,15^ (Table 2).

mGPD was also measured in a continuous spectrophotometric assay used by Madiraju et al^5^ in a reaction mixture of 10 mM Tris-chloride buffer pH 8.0, 50 mM KCl, 25 μM sodium azide, 1 mM EDTA, 50 μM cytochrome c with or without 25 mM L-glycerol phosphate as the DL mixture maintained at 550 nm at 38°. The background rate was recorded for 2 min before starting the enzyme rate by the addition of glycerol phosphate. When added, metformin was incubated along with the enzyme source in the reaction mixture for 10 min at 38° before measuring the background or enzyme rate as Madiraju et al^5^ did (Table 2).

mGPD activity when measured according to Rauchová et al^16^ was measured in a continuous spectrophotometric assay in a reaction mixture of 20 mM Tris-chloride buffer, pH 7.4, 50 mM KCl, 2 mM KCN, 1mM EDTA, 0.1 mM cytochrome c and with or without 25 mM L-glycerol phosphate as the DL mixture. After measuring a background rate for 2 min, the reaction was started by the addition of glycerol phosphate. The reaction rate was measured at 550 nm at 30°. When added, metformin was incubated with the enzyme source in the reaction mixture for 10 min at 30° before measuring the background or enzyme rates (Table 2).

Mitochondria were purified from liver, the INS-1 832/13 pancreatic beta cell line and the Huh 7.5 human liver cell line as previously described^17^.

Statistical significance of difference was calculated with Student’s t test.

## Acknowledgements

This work was supported by the Nowlin Family Trust of the InFaith Community Foundation.

## Author contributions

MJM designed the research and analyzed the data. IHA performed experiments and researched the data. MJL performed the experiments and researched the data. SWS performed experiments.

## Competing interests

Each author has no competing interest.

## REFERENCES

1. Madiraju, A., Erion, D., Rahimi, Y., Zhang, X.M., Braddock, D., Albright, R., Prigaro, B., Wood, J., Bhanot, S., MacDonald, M.J., Jurczak, M., Camporez, J.P., Lee, H-Y, Cline, G., Samuel, V., Kibbey, R.G., Shulman, G.I. (2014) Metformin suppresses gluconeogenesis by inhibiting mitochondrial glycerophosphate dehydrogenase. Nature 510:542–546.

2. MacDonald, M.J. (1981) High content of mitochondrial glyceral-3-phosphate dehydrogenase in pancreatic islets and its inhibition by diazoxide. J. Biol. Chem. 256:8287–8290.

3. Brown, L.J., Koza, R.A., Everett, C., Reitman, M.L., Marshall, L., Fahien, L.A., Kozak, L.P., MacDonald, M.J. (2002) Normal thyroid thermogenesis but reduced viability and adiposity in mice lacking the mitochondrial glycerol phosphate dehydrogenase. J. Biol. Chem. 277:32892–32898.

4. Eto, K., Tsubamoto, Y., Terauchi, Y., Sugiyama, T., Kishimoto, T., Takahashi, N., Yamauchi, N., Kubota, N., Murayama, S., Aizawa, T., Akanuma, Y., Aizawa, S., Kasai, H., Yazaki, Y., Kadowaki, T. (1999) Role of NADH shuttle system in glucose-induced activation of mitochondrial metabolism and insulin secretion. Science. 283:981–985.

5. Lardy, H.A., Paetkau, V., Walter, P. (1965) Paths of carbon in gluconeogenesis and lipogenesis: the role of mitochondria in supplying precursors of phosphoenolpyruvate. Proc. Natl. Acad. Sci. U.S.A. 53:1410–1415.

6. MacDonald, M.J. (1982) Evidence for the malate aspartate shuttle in pancreatic islets. Arch. Biochem. Biophys. 213:643–649.

7. MacDonald, M.J., Fahien, L.A., Brown, L.J., Hasan, N.M., Buss, J.D., Kendrick, M.A. (2005) Perspective: emerging evidence for signaling roles of mitochondrial anaplerotic products in insulin secretion. Am. J. Physiol. Endocrinol. Metab. 288:E1–15.

8. El-Mir, M.Y., Nogueira, V., Fontaine, E., Avéret, N., Rigoulet, M., Leverve, X. (2000) Dimethylbiguanide inhibits cell respiration via an indirect effect targeted on the respiratory chain complex I. J. Biol. Chem. 275:223–228.

9. Zhou, G., Myers, R., Li, Y., Chen, Y., Shen, X., Fenyk-Melody, J., Wu, M., Ventre, J., Doebber, T., Fujii, N., Musi, N., Hirshman, M.F., Goodyear, L.J., Moller, D.E. (2001) Role of AMP-activated protein kinase in mechanism of metformin action. J. Clin. Invest. 108:1167–1174.

10. Fryer, L.G., Parbu-Patel, A., Carling, D. (2002) The Anti-diabetic drugs rosiglitazone and metformin stimulate AMP-activated protein kinase through distinct signaling pathways. J. Biol. Chem. 277:25226–25232.

11. Shaw, R.J., Lamia, K.A., Vasquez, D., Koo, S.H., Bardeesy, N., Depinho, R.A., Montminy, M., Cantley, L.C. (2005) The kinase LKB1 mediates glucose homeostasis in liver and therapeutic effects of metformin. Science 310:1642–1646.

12. Misbin, R.I. (2004) The phantom of lactic acidosis due to metformin in patients with diabetes. Diabetes Care. 27:1791–1793.

13. Madiraju, A.K., Qiu, Y., Perry, R.J., Rahimi, Y., Zhang, X.M., Zhang, D., Camporez, J.G., Cline, G.W., Butrico, G.M., Kemp, B.E., Casals, G., Steinberg, G.R., Vatner, D.F., Petersen, K.F., Shulman, G.I. (2018) Metformin inhibits gluconeogenesis via a redox-dependent mechanism in vivo. Nat. Med. 24:1384–1394.

14. Gardner, R.S. (1974) A sensitive colorimetric assay for mitochondrial α-glycerophosphate dehydrogenase. Anal. Biochem. 59:272–276.

15. Dawson, A.P., Thorne, C.J.R. (1969) Preparation and some properties of l-3-glycerophosphate dehydrogenase from pig brain mitochondria. Biochem. J. 111:27–34.

16. Rauchová, H., Vokurková, M., Drahota, Z. (2014) Inhibition of mitochondrial glycerol-3-phosphate dehydrogenase by α-tocopheryl succinate. Int. J. Biochem. Cell Biol. 53:409–413.

17. Ansari, I.H., Longacre, M.J., Paulusma, C.C., Stoker, S.W., Kendrick, M.A., MacDonald, M.J. (2015) Characterization of P4 ATPase phospholipid translocases (flippases) in human and rat pancreatic beta cells: their gene silencing inhibits insulin secretion. J. Biol. Chem. 290:23110–23123

